# Phase Separation to Resolve Growth-Related Circuit Failures

**DOI:** 10.1101/2024.11.01.621586

**Authors:** Rong Zhang, Wangfei Yang, Rixin Zhang, Sadikshya Rijal, Abdelrahaman Youssef, Wenwei Zheng, Xiao-Jun Tian

## Abstract

Fluctuations in host cell growth poses a significant challenge to synthetic gene circuits, often disrupting circuit function. Existing solutions typically rely on circuit redesign with alternative topologies or additional control elements, yet a broadly applicable approach remains elusive. Here, we introduce a new strategy based on liquid-liquid phase separation (LLPS) to stabilize circuit performance. By engineering a self-activating circuit with transcription factors (TF) fused to an intrinsically disordered region (IDR), we enable the formation of TF condensates at the promoter region, maintaining local TF concentration despite growth-mediated dilution. This condensate formation preserves bistable memory in the self-activating circuit, demonstrating that phase separation can robustly counteract growth fluctuations, offering a novel design principle for resilient synthetic circuits.

## Introduction

Synthetic biology not only helps us understand the design principles of natural biological systems but also offers a powerful approach to constructing novel biological functions. However, feedback context factors, arising from the complex interplay between gene circuits and their host, pose significant challenges in designing robust synthetic gene circuits and predicting their dynamic behavior in response to variations in the host environment^1–4^. One key feedback factor is growth feedback, which stems from the reciprocal interactions between the gene circuit and the host cell’s growth rate^5–10^. The gene circuit draws on the host’s resources for gene expression, imposing a metabolic burden, while the growth rate influences resource availability and dilutes the circuit’s molecular concentration. Variations in growth conditions can significantly impair circuit functionality and, in some cases, lead to circuit failure^9–11^.

For example, Zhang et al found that the self-activation (SA) circuit is highly sensitive to cell growth^9^. The SA circuit, in theory, should operate as a bistable switch, maintaining a high expression ‘ON’ state once activated even after the initial signal is removed given its self-reinforcing properties. However, rapid cell growth in the fresh medium caused the circuit to quickly lose its memory, preventing it from returning to the ‘ON’ state, even after the host cells entered the stationary phase. The underlying issue was the dilution of transcription factors caused by rapid cell growth, which greatly reduced the transcriptional activity and compromised the circuit’s self-reinforcing capability.

Many control strategies have been proposed to address issues arising from circuit-host interactions^1,12–16^. Various feedback and feedforward controllers have been developed to minimize the interdependence between circuit genes caused by competition for limited cellular resources in the host cell^14–16^. For instance, Ceroni et al developed a CRISPR-dCas9-based negative feedback mechanism to alleviate cellular burden by adjusting the transcription of synthetic genes^12^. Barajas et al. introduced a feedforward controller that replenishes translational resources by modulating ppGpp levels upon synthetic gene induction, compensating for the metabolic burden and maintaining cellular function despite the increased load^14^. However, most of these strategies primarily focus on alleviating resource competition or optimizing resource reallocation. In addition, including more genes to implement these control mechanisms limits the scalability of these approaches.

In this work, we propose a simple yet effective engineering strategy to enhance the resilience of gene circuits to host cell’s division and growth by leveraging phase separation. As proof of concept, we show that the self-activation (SA) circuit becomes robust against growth-mediated dilution when an intrinsically disordered region (IDR) is fused to the circuit’s transcription factor (TF). This minimal modification enables the TF to form condensates at the promoter region, sustaining consistent transcriptional activity despite fluctuations in cellular growth and restoring the circuit’s memory maintenance capability. Our findings advance the design principles for building more stable and predictable gene circuits.

## Results

### Redesign of synthetic gene circuit using phase-separated condensate

To enhance the robustness of synthetic gene circuits against the effects of cell growth, we propose a redesign strategy that employs phase separation to concentrate transcription factors (TFs) at the promoter region where they are most needed (**Fig. 1a**). Specifically, we can achieve this by fusing an IDR to the TF, facilitating the formation of TF droplets or condensates at the promoter. As host cells grow and divide, while the average TF concentration may dilute, the size of the TF droplets may decrease, but the local TF concentration at the promoter remains effectively maintained.

**Figure 1.**
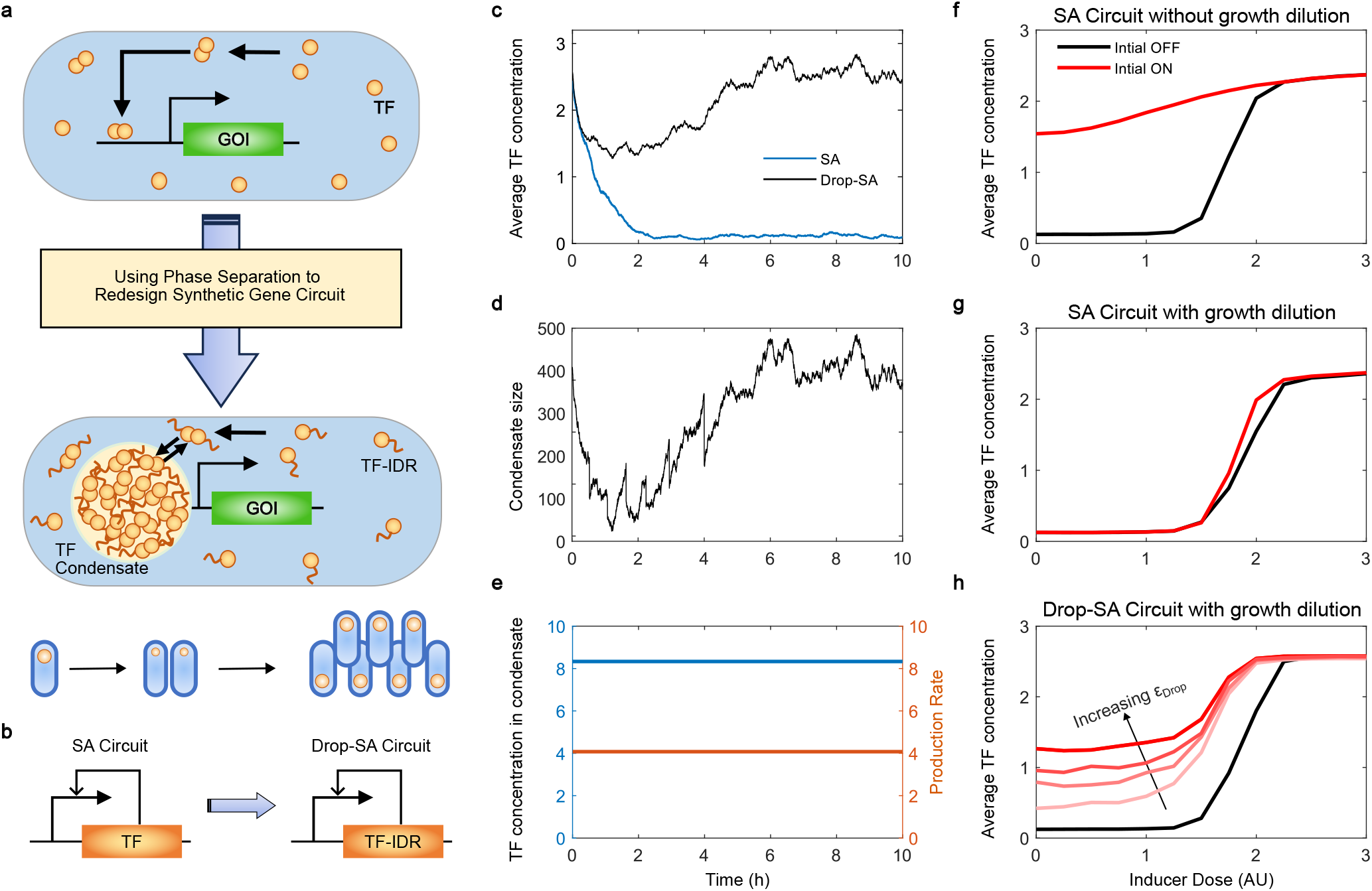
Theoretical analysis demonstrated redesigning synthetic gene circuits using phase separation could enhance circuit stability in response to host cell growth. (a) Schematic representation of the redesign of synthetic gene circuits using phase separation by fusing the transcription factor (TF) to an intrinsically disordered region (IDR), allowing TF droplets to form at the promoter region. (b) Application of the phase separation-based strategy to redesign the growth-sensitive self-activation (SA) circuit into the Drop-SA circuit. (c) Stochastic simulation results showing the dynamics of the average TF concentration in a single cell containing either the SA circuit (blue) or Drop-SA circuit (black) after diluting activated cells with high initial TF levels into fresh medium in batch culture. (d) The corresponding dynamics of droplet size in the cell containing the Drop-SA circuit as in (c). (e) Despite fluctuations in droplet size, the TF concentration within the droplet and the gene production rate remain stable throughout the dilution process. (f-h) Steady-state average TF concentrations of 1,000 simulated cells as a function of inducer dose, starting from an OFF (black) or ON (red) state. Results are shown for the SA circuit without growth dilution (f), the SA circuit with growth dilution (g), and the Drop-SA circuit with growth dilution under different strengths of effective concentration factor (εDrop) in the droplets (h).

The growth-sensitive SA circuit provides an excellent platform to study the impact of growth-mediated feedback on synthetic gene constructs. In this system, the dilution of transcription factor (TF) concentration due to cell growth directly leads to reduced transcriptional activity and memory loss. To counteract this growth-mediated memory loss, we employed this phase separation-based strategy to design an enhanced SA circuit, which we have termed the Droplet-Self-Activation (Drop-SA) circuit (**Fig. 1b**).

To first theoretically demonstrate that the Drop-SA circuit can counteract growth-mediated dilution effects, we developed a computational modeling framework to simulate the system’s dynamics. This framework incorporates stochastic gene expression using the Gillespie algorithm, cell growth, cell division, as well as condensate formation (see **Methods** for details). We compared the dynamics of average TF concentrations between the SA and Drop-SA systems after diluting activated cells with high initial TF levels (ON state) into fresh medium in batch culture. While the SA circuit rapidly loses its memory, transitioning to the OFF state where the TF concentration stays at basal level despite starting in the ON state, the Drop-SA circuit retains its memory and recovers to the original ON state, even though a moderate drop in TF concentration was observed (**Fig. 1c**). In the Drop-SA system, as cell growth and division cause a decline in the average TF concentration, the droplet size progressively shrinks from its initial larger state. However, the droplet persists and eventually recovers to its original size as the average TF concentration rebounds (**Fig. 1d**). Despite the significant fluctuations in condensate size, their persistent presence ensures a constant TF concentration within the condensates, supporting the consistent transcriptional activity (**Fig. 1e**).

We further compared the hysteresis properties of the SA and Drop-SA circuits by analyzing dose-response curves under two different initial conditions. Due to stochasticity, the results exhibited some level of cell-cell heterogeneity. For instance, under the same conditions, certain cells in the Drop-SA circuit experienced memory loss due to condensate dissipation (**Fig. S1a-c**). This cell-to-cell variability was further highlighted in **Fig. S1d-e**. To account for this variability, we conducted 1,000 stochastic simulations for each inducer dose and calculated the average TF concentration. In the absence of growth dilution, the SA circuit displayed a broad hysteresis range, with average TF concentrations dependent on initial conditions (**Fig. 1f**). However, under growth dilution conditions, the dose-response curves for the SA circuit no longer depended on initial conditions (**Fig. 1g**), indicating a loss of hysteresis due to dilution. Notably, the Drop-SA circuit successfully restored its hysteresis range with growth dilution to levels comparable to the SA circuit without growth dilution by strengthening the effective concentration factor (*ε_Drop_*) in the droplets (**Fig.1h**). Overall, theoretical analysis demonstrates that phase separation prevents memory loss in the SA circuit, making its function more resilient to host cell growth.

### Fusing the circuit’s transcriptional factor with an intrinsically disordered region (IDR) to facilitate its condensate formation

Intrinsically disordered regions (IDRs) promote phase separation by facilitating flexible, multivalent interactions that drive the dynamic assembly of biomolecular condensates^17–19^. A well-known example is the N-terminal domain of the Fused in Sarcoma (FUS) protein (FUSn), a natural IDR that has been extensively studied for its ability to promote liquid-like condensate formation through these multivalent interactions^20–23^. Additionally, synthetic IDRs, such as resilin-like polypeptides (RLPs), have been engineered to drive biomolecular condensate formation with tunable properties^24–26^. Both FUSn and RLPs exhibit phase separation behavior characterized by the upper critical solution temperature (UCST) mechanism, meaning they tend to form condensates at lower temperatures.

We first constructed one bicistronic SA circuit **OP174** (**Fig. 2a**), where the transcriptional factor AraC and the reporter green fluorescent protein (GFP) are transcribed into a single mRNA strand under the same promoter Pbad but translated separately with each of its ribosome binding site (RBS). In the presence of inducer L-arabinose (Lara), AraC forms a dimer and binds to the promoter, driving the expression of both AraC and GFP. The GFP expressed from this circuit does not form condensates at any expression level in *E. coli* (**Fig. 2b**). To validate whether FUSn and RLP20 facilitate condensate formation, we constructed two gene circuits, **CT123 and CT124**, by fusing RLP20 or FUSn to the C-terminus of GFP (**Fig. S2**). We observed that both GFP-FUSn and GFP-RLP20 form small, intensely fluorescent droplets at the polar regions of *E. coli*, with diffuse weaker fluorescence dispersed throughout the surrounding dilute phase (**Fig. 2c-d**).

**Figure 2.**
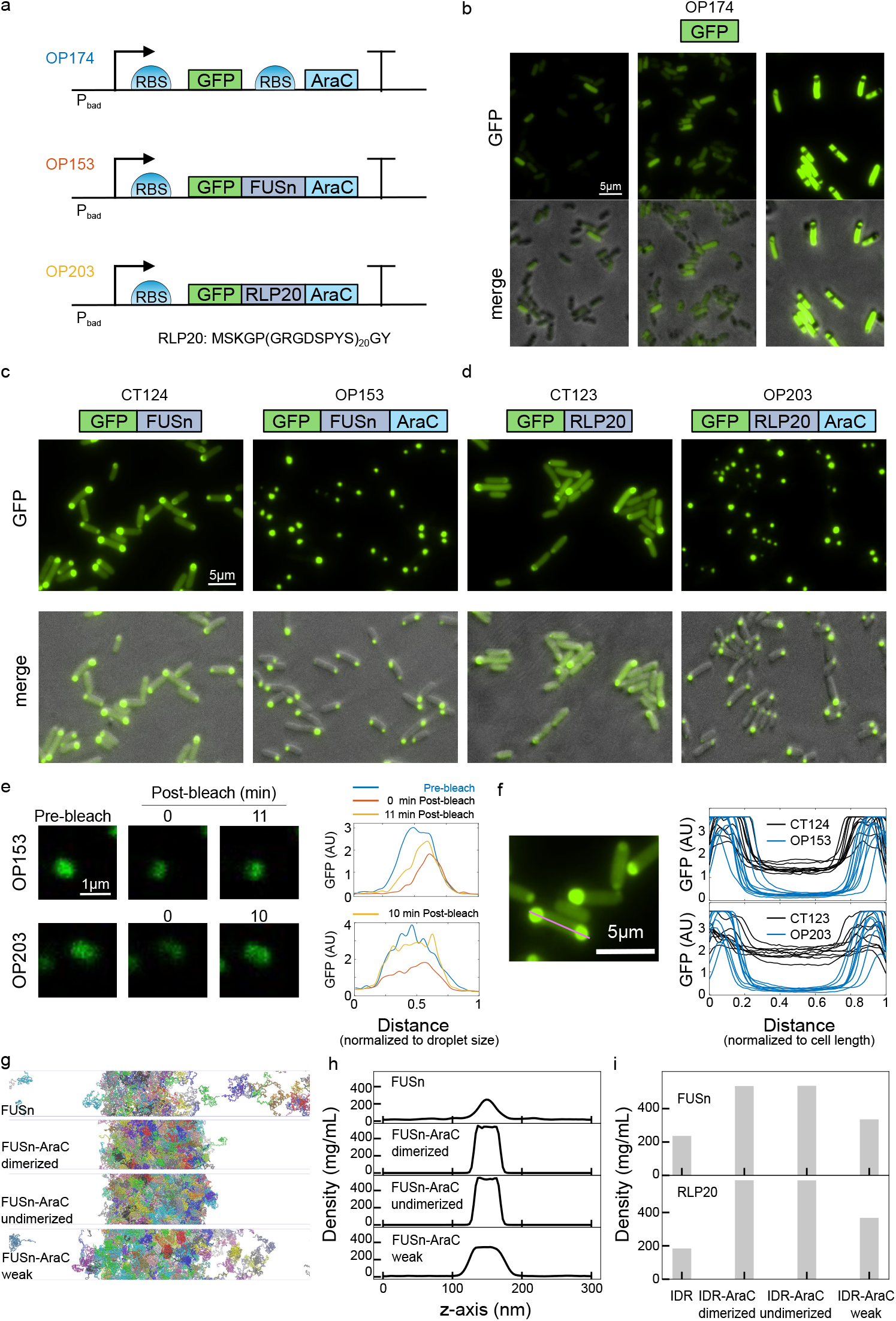
Fusing Intrinsically Disordered Regions (IDRs) to Transcription Factors Promotes its Condensate Formation. (a) Schematic of the three self-activation (SA) circuits: OP174, OP153, and OP203. In OP174, the transcription factor AraC and the reporter green fluorescent protein (GFP) are co-expressed bicistronically under the promoter Pbad. In OP153 and OP203, fusion proteins GFP-FUSn-AraC and GFP-RLP20-AraC are expressed monocistronically under promoter Pbad. (b) Fluorescence microscopy images of cells containing circuit OP174, showing the scattered distribution of GFP after overnight induction with varying concentrations of L-arabinose (Lara): 0.625 ppm, 2.5 ppm, and 25 ppm. (c-d) Fluorescence microscopy images showing intracellular droplet formation of GFP-FUSn (circuit CT124), GFP-FUSn-AraC (circuit OP153), GFP-RLP20 (circuit CT123), and GFP-RLP20-AraC (circuit OP203) at (c) 4 hours and (d) 4.5 hours post-induction with Lara at 37°C. CT123 and CT124 were induced with 12.5 ppm Lara, while OP153 and OP203 were induced with 50 ppm Lara to achieve similar levels of total GFP intensities. (e) Representative fluorescence recovery after photobleaching (FRAP) images and corresponding quantified data for single droplets in *E. coli* expressing GFP-FUSn-AraC (OP153) or GFP-RLP20-AraC (OP203) fusion proteins, showing fluorescence levels pre-bleach, immediately post-bleach (0 minutes), and at 10- or 11-minutes post-bleach. (f) Line-profile of fluorescence intensity across single *E. coli* cells (indicated by the magenta line) expressing GFP-FUSn (CT124), GFP-FUSn-AraC (OP153), GFP-RLP20 (CT123), and GFP-RLP20-AraC (OP203), demonstrating the relative concentration difference of fusion proteins between the droplet phase and the surrounding cytoplasm (dilution phase). Each curve represents one single cell. (g) Slab simulations illustrating the phase coexistence in systems with FUSn alone, or with FUSn-AraC fusion protein, considering three cases for the latter: strong AraC dimerization, no AraC dimerization, and no AraC dimerization with weakened AraC interactions. (h) Protein density profiles corresponding to the slab simulations shown in (g). (i) Protein densities within the condensate phase from the slab simulations for both FUSn and RLP20 systems.

We then constructed two Drop-SA circuits, **OP153 and OP203**, by further fusing AraC to the C-terminus of GFP-FUSn and GFP-RLP20 (**Fig. 2a**). Similarly, small droplets were observed at the polar regions of cells. To verify that the condensates result from phase separation, we performed Fluorescence Recovery After Photobleaching (FRAP). After photobleaching a specific region of the droplets in cells containing circuits OP153 or OP203, we monitored the recovery of fluorescence over time. The fluorescence exhibited rapid recovery, reaching a plateau approximately 10 to 11 minutes post-bleach. This observation indicates a dynamic exchange of molecules within the droplets and provides confirmation of liquid-liquid phase separation (LLPS) (**Fig. 2e)**. One notable feature in OP153 and OP203 systems is that the fluorescence in the dilute phase was barely detectable. That is, the concentration difference between the droplet phase and the dilute phase of GFP-FUSn-AraC and GFP-RLP20-AraC is significantly enhanced compared with GFP-FUSn and GFP-RLP20 (**Fig. 2f**). To investigate the underlying mechanism, we conducted in silico coarse-grained molecular dynamics simulations using a hybrid model (see **Methods** for details).

Simulations of IDP alone (FUSn and RLP20) and fusion proteins (FUSn-AraC and RLP20-AraC) using our slab sampling strategy reveals the phase coexistence in all systems (**Fig. 2g and S3a**). To assess the phase separation tendencies, we calculated the density profiles, which display a condensate phase in the box center and a dilute phase at the edges (**Fig. 2h and S3b**). The enhanced protein concentration in the condensate phase (**Fig. 2i)** confirms that attaching AraC to either FUSn or RLP20 improves phase separation. To investigate whether this enhancement arises from AraC’s general multivalent interactions or specific dimerization, we gradually reduced the intermolecular dimerization interactions between AraC monomers, producing systems with varying dimer fractions in the condensate (**Fig. S3c-d**). Interestingly, the concentration within the condensate phase remained almost constant across all dimer fractions, indicating that AraC’s dimerization is not the primary driver of enhanced condensate formation. We also examined systems with reduced hydrophobic interactions between AraC’s folded domains and other components, entirely omitting AraC’s dimerization. As shown in **Fig. 2i**, the protein density in the condensate phase decreased as multivalent interaction weakened, suggesting that AraC enhances phase separation mainly through general multivalent interactions, rather than dimerization. This mechanism aligns with similar observations in other systems ^27,28^.

### Phase separation enhances gene expression and accelerates activation dynamics of self-activation circuits

We further investigated how phase separation of transcription factors affects gene expression levels. To do this, we constructed three open-loop gene circuits, CT115, CT116, and CT118, in which AraC was expressed either alone or fused to the N- or C-terminus of FUSn under a constitutive promoter. The reporter gene GFP was placed under the control of the Pbad promoter (**Fig. S4a**). GFP expression levels were measured 16 hours after treatment with varying doses of L-arabinose using a plate reader. The dose-response curve revealed that fusing FUSn to the N-terminus of AraC enhanced gene expression by 2-3 folds while fusing it to the C-terminus of AraC resulted in minimal changes to expression levels (**Fig. S4b**). This is consistent with previous findings ^26,29^. In addition, we constructed three closed-loop SA circuits, **OP174, OP175, and OP176**, in which AraC alone, fused to the N- or C-terminus of FUSn, was also placed under the control of the Pbad promoter (**Fig. S5a**). The dose-response curves from these circuits confirmed that fusing FUSn to the N-terminus of AraC significantly enhances gene expression compared to fusing to the C-terminus (**Fig. S5b**).

We further tested whether the inclusion of an IDR domain would enhance gene expression in circuits OP153 and OP203, where the tri-fusion proteins GFP-FUSn-AraC and GFP-RLP20-AraC are produced. As a control, we constructed a similar circuit (OP177) using a GS linker to replace the IDR. Based on the dose-response curves (**Fig. 3a**), we observed that the fold change in GFP levels was significantly higher in circuits OP153 and OP203 compared to the control circuit OP177. Taken together, phase-separated transcriptional factors could amplify gene expression.

**Figure 3.**
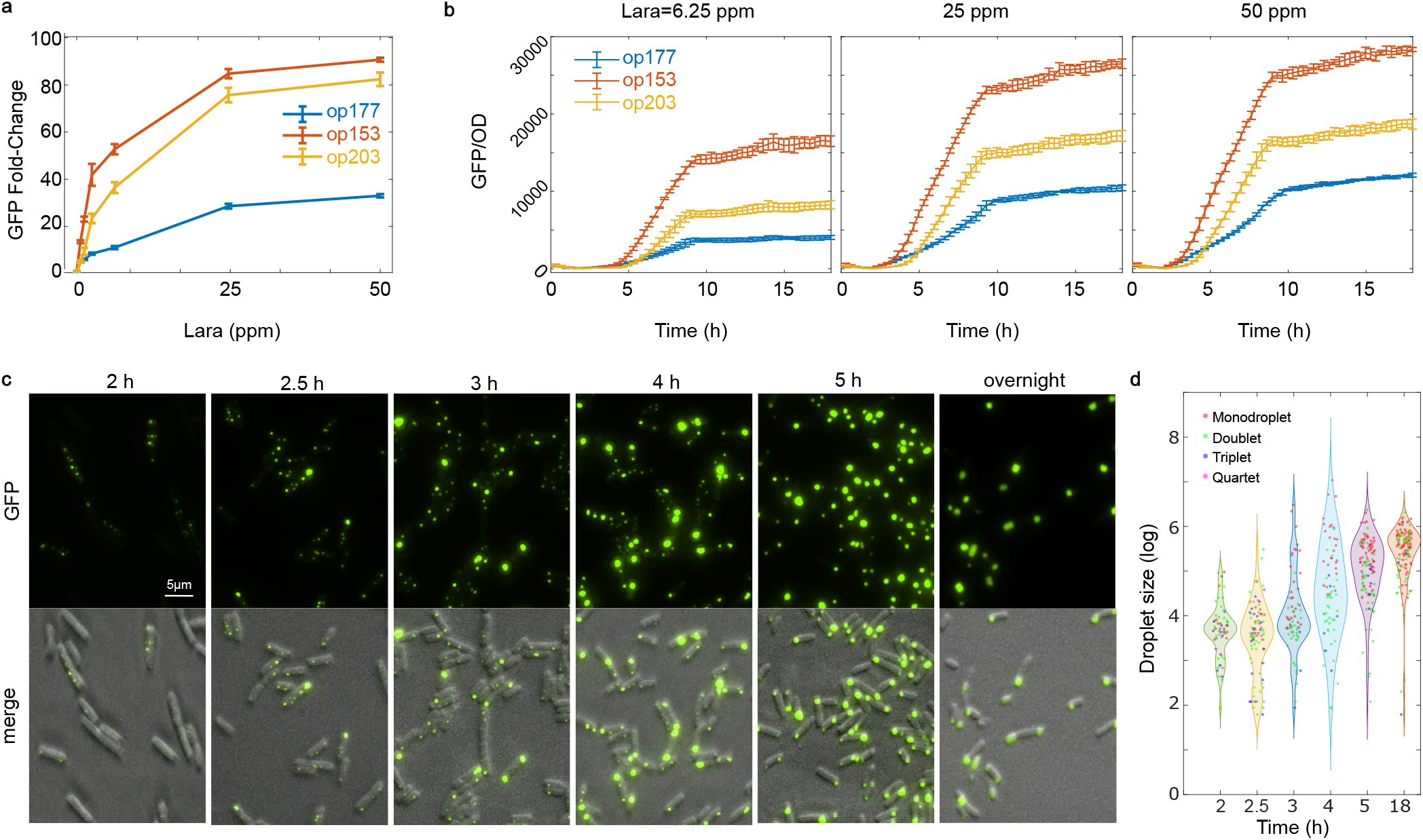
Phase separation enhances gene expression and accelerates activation dynamics of self-activation circuits. (a) Dose-response curves showing the dependence of GFP fold-changes on inducer concentration for circuits OP177 (GFP-GSlinker-AraC), OP153 (GFP-FUSn-AraC), and OP203 (GFP-RLP20-AraC). Data in the curve represents means ± s.d. n=3. (b) Time-course analysis of GFP/OD levels in cells expressing GFP-GSlinker-AraC (OP177), GFP-FUSn-AraC (OP153), and GFP-RLP20-AraC (OP203), starting from the OFF state, with varying L-arabinose (Lara) concentrations: 0.625 ppm, 2.5 ppm, and 25 ppm. Data in the curve represents means ± s.d. n=3. (c) Fluorescence microscopy images showing the dynamics of droplet formation in cells induced to express GFP-FUSn-AraC (OP153) for 2, 2.5, 3, 4, and 5 hours, and overnight with 50 ppm Lara at 34°C. (d) Distribution of quantified droplet size and number in single cells at different time points. Each dot represents one single droplet.

We further measured the dynamics of gene expression under various inducer doses. We found that the switch activation was accelerated in circuits OP153 and OP203 compared to the control circuit OP177 under the same inducer doses (**Fig. 3b**). **Fig. 3c** and **Fig. S6a** show the dynamics of droplet formation in the Drop-SA circuits OP153 and OP203. We observed the formation of small droplets in each cell at 2∼2.5 hours post-induction, reflecting the time required for transcription, translation, and the accumulation of sufficient TF levels to initiate droplet formation. The number of droplets per cell varied significantly, ranging from one to four, with more than four droplets being rare. Over time, the droplets grew larger as fused proteins, GFP-FUSn-AraC and GFP-RLP20-AraC, accumulated, with smaller droplets merging to form larger ones. By overnight incubation, most cells contained one or two large droplets, predominantly localized at the poles. To see clearly, we quantified the droplet number and size in each cell. As shown in **Fig. 3d and Fig.S6b**, while monodroplets, doublets, triplets, and quartets were common at earlier time points, their sizes generally increased, and fewer triplet and quartet droplets were observed later. Ultimately, only monodroplets or doublets, larger in size, remained at later time points. Interestingly, in cells containing the OP177 circuit, the fusion protein GFP-GSlinker-AraC displayed an uneven distribution, with irregularly shaped fluorescent patches appearing during the early hours (**Fig. S7**). After 5 hours, condensates formed at the cell poles, likely due to significant concentration inhomogeneity of the transcription factor (TF), driven by binding to its target in the plasmids and exclusion from the host chromosome^30,31^. To confirm that the uneven TF spatial distribution is not caused by the GS linker, we directly fused GFP and AraC. The GFP-AraC fusion protein exhibited the same dynamic behavior (**Fig. S8**). In summary, fusing an IDR domain can significantly enhance gene expression and accelerate the activation of the genetic switch.

### Phase-separated condensates enhance the memory retention of self-activation circuits

To assess whether memory could be retained in the Drop-SA circuits, we first activated circuits OP177, OP153, and OP203 overnight using a high dose of Lara (Lara=50 ppm) and then diluted the cells into fresh medium containing a lower dose (Lara=2.5 ppm). We captured images of the cells at different time points from 0 to 8 hours post-dilution. As shown in **Fig. 4a-c**, initially, most cells across all three circuits displayed 1 or 2 large droplets at the polar regions of *E. coli*. As the cells grew and divided over time, the droplet sizes decreased, and more than two smaller droplets appeared in some cells. For OP177, most droplets began disappearing in most cells by 3-4 hours, whereas small droplets persisted in the OP153 and OP203 systems as shown in **Fig. S9** with intensified fluorescence. Droplets in OP153 and OP203 began regrowing to larger sizes between 5-6 hours, while no droplet regrowth was observed in OP177. By 8 hours, the droplets had recovered to their original size in most cells for OP153 and OP203, but only in a few cells in the OP177 system. This was also confirmed in quantified average GFP per cell (**Fig. S10**). This strongly suggests that fusing the IDR domain with the transcriptional factors in the SA circuit enhances memory retention.

**Figure 4.**
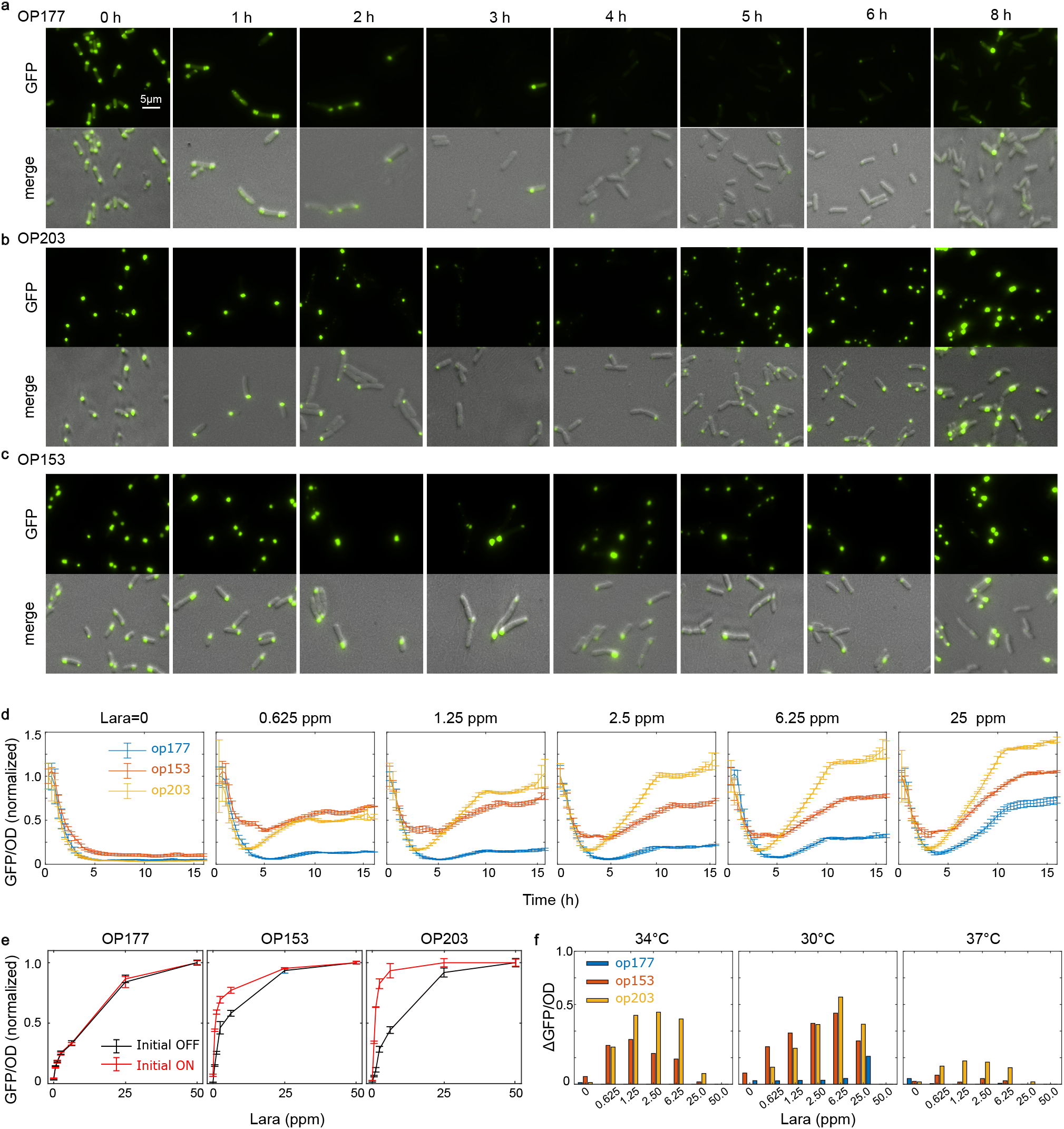
Phase-separated condensates enhance the memory retention of self-activation circuits. (a-c) Fluorescence microscopy images showing the dynamics of droplet shrinkage due to cell growth and subsequent recovery in cells containing circuits OP203 and OP153 at 34°C. (d) Time-course analysis of GFP/OD levels after diluting activated cells with high GFP levels into fresh medium containing varying concentrations of L-arabinose (0 to 25 ppm) at 34°C. Data in the curve represent means ± s.d. n=3. (e) Dose-response curves for steady-state GFP/OD levels in cells containing circuits OP177 (left), OP153 (middle), and OP203 (right), starting from either OFF or ON states at 34°C. Data in the curve represent means ± s.d. n=3. (f) The difference in steady-state GFP/OD levels between starting from ON and OFF states (ΔGFP/OD) in each system at different temperatures: 34°C (left), 30°C (middle), and 37°C (right). Data in the curve represent means. n=3.

We also systematically measured the GFP dynamics for all three systems after diluting the activated cells into fresh medium containing different lower concentrations of Lara (0–25 ppm) using a plate reader. As shown in **Fig. 4d**, the average GFP levels in all three systems rapidly decreased to low levels without recovery in the medium lacking Lara. However, in the OP153 and OP203 systems, the GFP levels did not drop as low as it did in OP177 system. By 4–5 hours, they began to recover, eventually reaching different higher levels depending on the Lara concentration. We evaluated the hysteresis range by comparing the dose-response curves with ON and OFF initial conditions. As shown in **Fig. 4e**, no hysteresis was observed in the OP177 circuit, as the two curves were nearly identical, indicating that the system reached the same steady state regardless of its initial condition.

This suggests that memory was lost due to growth-mediated dilution. In contrast, both the OP153 and OP203 circuits displayed a significant hysteresis, where GFP levels were higher with an ON initial condition than with an OFF one. We quantified the hysteresis by calculating the difference between the two dose-response curves. As shown in **Fig. 4f**, this difference is significantly higher in circuits OP153 and OP203 compared to the circuit OP177.

Further experiments revealed temperature dependence, where the hysteresis became more pronounced at 30°C and diminished at 37°C (**Fig. S11, Fig. 4f**). One reason for this is that the slower growth rate at lower temperatures, which reduces dilution and provides more time for the system to recover (**Fig. S12**). Additionally, both FUSn and RLP20 exhibit upper critical solution temperature (UCST) behavior, meaning that lower temperatures favor condensate formation. As a result, at lower temperatures, the condensates in the OP153 and OP203 systems are less likely to dissolve during rapid cell growth and division, enhancing the stability of the circuit. In addition, we evaluated the hysteresis of the three systems under varying nutrient levels. As shown in **Fig. S13**, hysteresis was enhanced in all three systems at lower nutrient levels, as slower growth rates and limited growth times reduce dilution effects, but the OP153 and OP203 circuits continued to exhibit much more hysteresis ranges compared to OP177. Taken together, redesigning the SA circuit to incorporate phase separation greatly enhanced its bistable function, making it more robust.

## Discussion

We proposed a simple yet elegant approach to enhance the robustness of a synthetic gene circuit against host cell growth by making minimal modifications. By fusing an IDR to the circuit’s transcription factor, we enabled it to concentrate at the promoter region, ensuring stable transcription even under rapid dilution of the average transcription factor concentration caused by fast host cell growth. We validated this concept by demonstrating that introducing phase separation properties to the growth-sensitive SA circuit allowed it to retain its memory in response to growth dilution, thereby preserving its functionality under challenging conditions.

Numerous strategies have been employed to alleviate metabolic burden and growth-mediated effects. For example, Ceroni et al implemented a dCas9-based negative feedback to control gene expression in synthetic gene circuits in response to metabolic burden^12^. Barajas et al employed a feedforward controller to modulate growth rate and compensate for the metabolic burden of exogenous gene expression^14^. Stone et al introduced a straightforward supplementary repressive regulation within the circuit to enhance the circuit stability under growth feedback^15^. Unlike labor-intensive approaches that involve redesigning gene circuits with complex topologies or integrating additional genes for feedback or feedforward control, phase separation provides a highly generalizable and efficient solution. Simply fusing an IDR into the original circuit design enables easy implementation with minimal modifications. This approach can be seamlessly applied to other systems without imposing a significant cellular burden or consuming excessive resources.

Phase separation serves as a mechanism for spatial organization and functional compartmentalization within cells, playing an important role in biological signal processing and regulation^17,32^. For instance, stress granules form to sequester untranslated mRNA and protect it from degradation under stress conditions^33,34^. Additionally, phase separation also can act as a noise reduction mechanism by insulating the protein concentration in the dilute phase from changes in total concentration^35^. It is important to note that the rapid cell growth during the exponential phase causes substantial dilution of transcription factor concentration, representing a large fluctuation that can easily switch off the SA circuit^9^. This effect arises from the circuit’s strong dependence on high TF concentration to sustain activation, making the transcription rate highly sensitive to changes in TF concentration within the self-activation system. Nevertheless, phase-separating transcription factors can mitigate concentration drops by maintaining relatively stable transcription levels.

Here we provided proof-of-concept for using phase separation to design gene circuits that are robust to host cell growth rate variations. Refining various parameters, such as adjusting the concentration threshold required for phase separation and modulating the diffusion rate of the protein molecules between the droplet and dilute phases, could effectively prolong the stability of the condensate and mitigate memory loss due to dilution. Our study primarily focused on condensing transcription factors using IDR at the promoter region. Exploring the potential integration of phase separation into the translational process could offer further optimization avenues for this strategy. Future work will exploit orthogonal or multicomponent condensate to scale up synthetic gene circuit design, addressing challenges such as resource competition between circuit modules^36–38^. Harnessing phase separation for sophisticated and robust synthetic gene circuits presents both opportunities and challenges. While it provides some advantages such as compartmentalization and concentration buffering, it also introduces complexity to the system. For example, unlike in single-component condensate, the dilute and dense concentrations might be unfixed in multicomponent systems^39–41^. Balancing these variables will be crucial for optimizing the performance of synthetic gene circuits and fully exploiting the advantages of phase separation in gene circuit design.

## Data availability

Source data are available online.

## Code availability

The source scripts and code for simulation are also available at https://github.com/TianLab-ASU/PhaseSeparation_Growth_CircuitMemory and https://github.com/wzhenglab/AraC-2024.git.

## Acknowledgments

The authors thank Lingchong You and Ashutosh Chilkoti for providing the RLP20 plasmid. This work was supported by a grant from the US National Institutes of Health (R35GM142896 to X.-J.T., and R35GM146814 to W.Z.).

## Author Contributions

X.-J.T. conceived the study. Ro.Z., X.-J.T. designed the experiments and interpreted the data. Ro.Z., Ri.Z., S. R., A. Y., performed the experiments. Ro.Z., Ri.Z., X.-J.T. analyzed the data. W.Y., W. Z., X.-J.T. performed the in-silico simulation and analysis. R.Z., W.Y., W. Z., X.-J.T. wrote the paper.

## References

1. Stone, A., Youssef, A., Rijal, S., Zhang, R. & Tian, X.-J. Context-dependent redesign of robust synthetic gene circuits. Trends Biotechnol. 42, 895–909 (2024).

2. Del Vecchio, D. Modularity, context-dependence, and insulation in engineered biological circuits. Trends Biotechnol. 33, 111–119 (2015).

3. Borkowski, O., Ceroni, F., Stan, G.-B. & Ellis, T. Overloaded and stressed: whole-cell considerations for bacterial synthetic biology. Curr. Opin. Microbiol. 33, 123–130 (2016).

4. Boo, A., Ellis, T. & Stan, G.-B. Host-aware synthetic biology. Curr. Opin. Syst. Biol. 14, 66–72 (2019).

5. Klumpp, S. & Hwa, T. Bacterial growth: global effects on gene expression, growth feedback and proteome partition. Curr. Opin. Biotechnol. 28, 96–102 (2014).

6. Ceroni, F. et al. Quantifying cellular capacity identifies gene expression designs with reduced burden. Nat. Methods 12, 415–418 (2015).

7. Liao, C., Blanchard, A. E. & Lu, T. An integrative circuit–host modelling framework for predicting synthetic gene network behaviours. Nat. Microbiol. 2, 1658–1666 (2017).

8. Tan, C., Marguet, P. & You, L. Emergent bistability by a growth-modulating positive feedback circuit. Nat. Chem. Biol. 5, 842–848 (2009).

9. Zhang, R. et al. Topology-dependent interference of synthetic gene circuit function by growth feedback. Nat. Chem. Biol. 16, 695–701 (2020).

10. Nikolados, E.-M., Weiße, A. Y., Ceroni, F. & Oyarzún, D. A. Growth Defects and Loss-of-Function in Synthetic Gene Circuits. ACS Synth. Biol. 8, 1231–1240 (2019).

11. Kong, L.-W., Shi, W., Tian, X.-J. & Lai, Y.-C. Effects of growth feedback on gene circuits: A dynamical understanding. eLife 12, (2023).

12. Ceroni, F. et al. Burden-driven feedback control of gene expression. Nat. Methods 15, 387–393 (2018).

13. Barajas, C. & Del Vecchio, D. Synthetic biology by controller design. Curr. Opin. Biotechnol. 78, 102837 (2022).

14. Barajas, C., Huang, H.-H., Gibson, J., Sandoval, L. & Del Vecchio, D. Feedforward growth rate control mitigates gene activation burden. Nat. Commun. 13, 7054 (2022).

15. Stone, A., Rijal, S., Zhang, R. & Tian, X.-J. Enhancing circuit stability under growth feedback with supplementary repressive regulation. Nucleic Acids Res. gkad1233 (2024).

16. Jones, R. D. et al. An endoribonuclease-based feedforward controller for decoupling resource-limited genetic modules in mammalian cells. Nat Commun 11, 5690 (2020).

17. Lyon, A. S., Peeples, W. B. & Rosen, M. K. A framework for understanding the functions of biomolecular condensates across scales. Nat. Rev. Mol. Cell Biol. 22, 215–235 (2021).

18. Alberti, S., Gladfelter, A. & Mittag, T. Considerations and Challenges in Studying Liquid-Liquid Phase Separation and Biomolecular Condensates. Cell 176, 419–434 (2019).

19. Dai, Y., You, L. & Chilkoti, A. Engineering synthetic biomolecular condensates. Nat. Rev. Bioeng. 1–15 (2023) doi:10.1038/s44222-023-00052-6.

20. Zhao, E. M. et al. Light-based control of metabolic flux through assembly of synthetic organelles. Nat. Chem. Biol. 15, 589–597 (2019).

21. Murakami, T. et al. ALS/FTD Mutation-Induced Phase Transition of FUS Liquid Droplets and Reversible Hydrogels into Irreversible Hydrogels Impairs RNP Granule Function. Neuron 88, 678–690 (2015).

22. Bremer, A. et al. Deciphering how naturally occurring sequence features impact the phase behaviours of disordered prion-like domains. Nat. Chem. 14, 196–207 (2022).

23. Patel, A. et al. A Liquid-to-Solid Phase Transition of the ALS Protein FUS Accelerated by Disease Mutation. Cell 162, 1066–1077 (2015).

24. Dzuricky, M., Rogers, B. A., Shahid, A., Cremer, P. S. & Chilkoti, A. De novo engineering of intracellular condensates using artificial disordered proteins. Nat. Chem. 12, 814–825 (2020).

25. Quiroz, F. G. & Chilkoti, A. Sequence heuristics to encode phase behaviour in intrinsically disordered protein polymers. Nat. Mater. 14, 1164–1171 (2015).

26. Dai, Y. et al. Programmable synthetic biomolecular condensates for cellular control. Nat. Chem. Biol. 1– 11 (2023) doi:10.1038/s41589-022-01252-8.

27. Elbaum-Garfinkle, S. et al. The disordered P granule protein LAF-1 drives phase separation into droplets with tunable viscosity and dynamics. Proc. Natl. Acad. Sci. 112, 7189–7194 (2015).

28. Dignon, G. L., Zheng, W., Kim, Y. C., Best, R. B. & Mittal, J. Sequence determinants of protein phase behavior from a coarse-grained model. PLoS Comput. Biol. 14, e1005941 (2018).

29. Schneider, N. et al. Liquid-liquid phase separation of light-inducible transcription factors increases transcription activation in mammalian cells and mice. Sci. Adv. 7, eabd3568 (2021).

30. Kuhlman, T. E. & Cox, E. C. Gene location and DNA density determine transcription factor distributions in Escherichia coli. Mol. Syst. Biol. 8, 610 (2012).

31. Kuhlman, T. E. & Cox, E. C. DNA-binding-protein inhomogeneity in $E.$ coli modeled as biphasic facilitated diffusion. Phys. Rev. E 88, 022701 (2013).

32. Holehouse, A. S. & Pappu, R. V. Functional Implications of Intracellular Phase Transitions. Biochemistry 57, 2415–2423 (2018).

33. Anderson, P. & Kedersha, N. Stress granules: the Tao of RNA triage. Trends Biochem. Sci. 33, 141–150 (2008).

34. Riback, J. A. et al. Stress-Triggered Phase Separation Is an Adaptive, Evolutionarily Tuned Response. Cell 168, 1028–1040.e19 (2017).

35. Klosin, A. et al. Phase separation provides a mechanism to reduce noise in cells. Science 367, 464–468 (2020).

36. Zhang, R. et al. Winner-takes-all resource competition redirects cascading cell fate transitions. Nat. Commun. 12, 853 (2021).

37. Goetz, H., Stone, A., Zhang, R., Lai, Y. & Tian, X. Double-Edged Role of Resource Competition in Gene Expression Noise and Control. Adv. Genet. 3, 2100050 (2022).

38. Gyorgy, A. et al. Isocost Lines Describe the Cellular Economy of Genetic Circuits. Biophys. J. 109, 639– 646 (2015).

39. Riback, J. A. et al. Composition-dependent thermodynamics of intracellular phase separation. Nature 581, 209–214 (2020).

40. Deviri, D. & Safran, S. A. Physical theory of biological noise buffering by multicomponent phase separation. Proc. Natl. Acad. Sci. 118, e2100099118 (2021).

41. Dai, Y. et al. Biomolecular condensates regulate cellular electrochemical equilibria. Cell 187, 5951–5966.e18 (2024).

42. Tesei, G. & Lindorff-Larsen, K. Improved predictions of phase behaviour of intrinsically disordered proteins by tuning the interaction range. Open Res. Eur. 2, 94 (2022).

43. Soisson, S. M., MacDougall-Shackleton, B., Schleif, R. & Wolberger, C. Structural basis for ligand-regulated oligomerization of AraC. Science 276, 421–425 (1997).

44. Rodgers, M. E. & Schleif, R. Solution structure of the DNA binding domain of AraC protein. Proteins 77, 202–208 (2009).

45. Eastman, P. et al. OpenMM 7: Rapid development of high performance algorithms for molecular dynamics. PLoS Comput. Biol. 13, e1005659 (2017).

46. Szatmári, D. et al. Intracellular ion concentrations and cation-dependent remodelling of bacterial MreB assemblies. Sci. Rep. 10, 12002 (2020).

