## Supplementary Figure 1-13 for "Phase Separation to Resolve Growth-Related Circuit Failures"

### Supplementary Figures

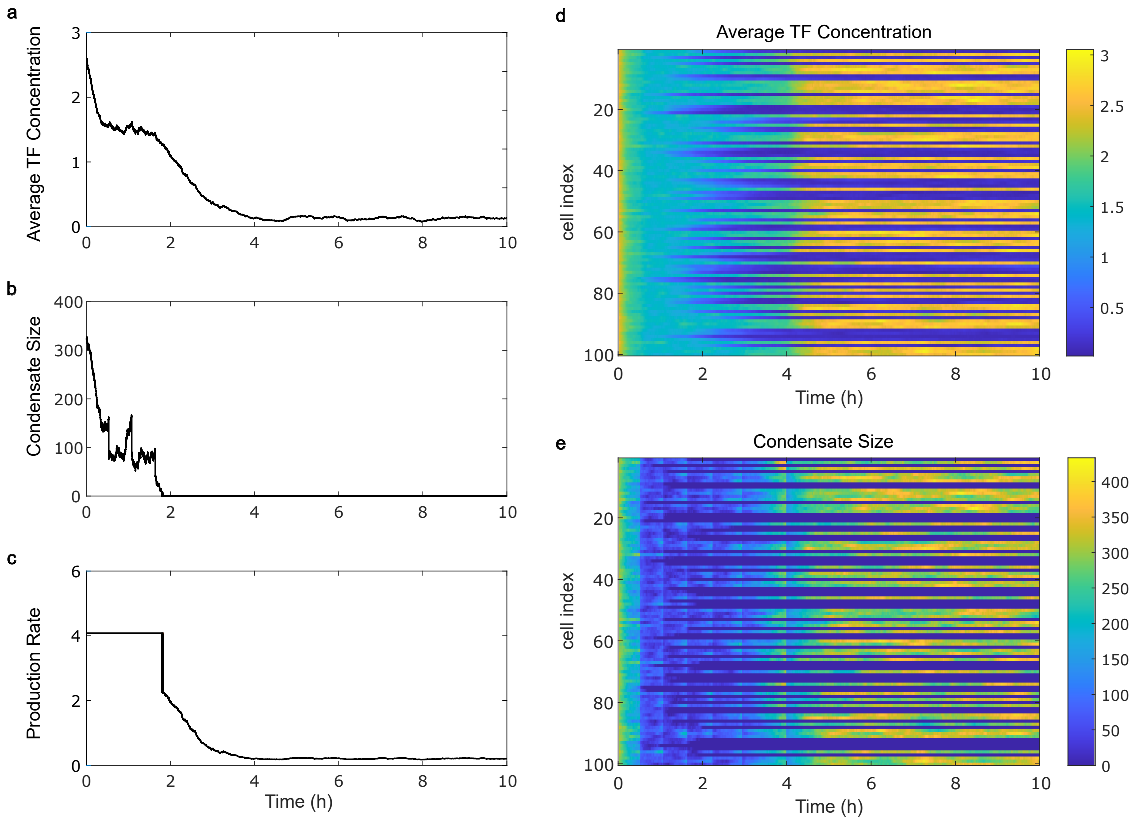


**Supplementary figure 1. Stochasticity and Cell-to-Cell Heterogeneity in Memory Retention of the Drop-SA Circuit.** (a-c) Stochastic simulation results illustrating the dynamics of a Drop-SA circuit in a cell that lost memory after dilution into fresh medium in batch culture, despite starting with high initial TF levels. The panels show (a) the average TF concentration, (b) the droplet size, and (c) the gene production rate over time. (d-e) Dynamics of the average TF concentration (d) and droplet size (e) across 100 simulated cells, highlighting the cell-to-cell heterogeneity in memory retention.

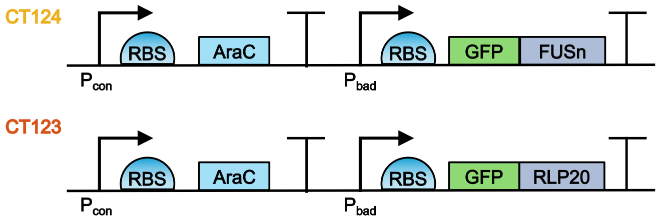


**Supplementary figure 2.** Schematic illustrations of the CT123 and CT124 circuit designs. The TF AraC is constitutively expressed, while the fusion proteins are placed under Pbad promoter.

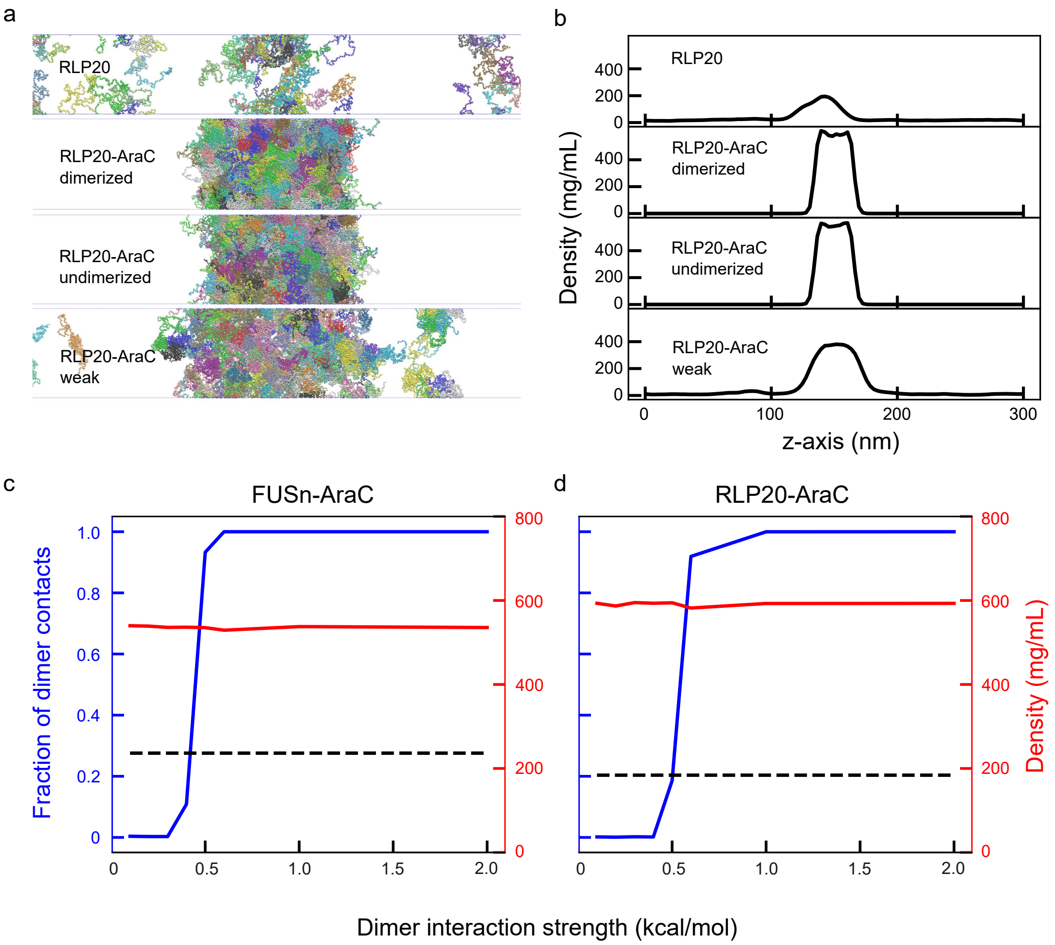


**Supplementary figure 3. Additional data for coarse-grained (CG) molecular dynamics simulations of FUSn and RLP variants.** (a) Slab simulations illustrating the phase coexistence in systems with RLP20 alone, or with RLP20-AraC fusion protein, considering three cases for the latter: strong AraC dimerization, no AraC dimerization, and no AraC dimerization with weakened AraC interactions. (b) Protein density profiles corresponding to the slab simulations shown in (a). (c-d) Protein densities within the condensate phase (red curve) and fraction of the dimer contacts (blue curve) as a function of dimer interaction strength in systems with FUSn-AarC (c) or RLP-AraC (d). Black dashed lines indicate the protein densities within the condensate phase for the systems with the corresponding IDR alone.

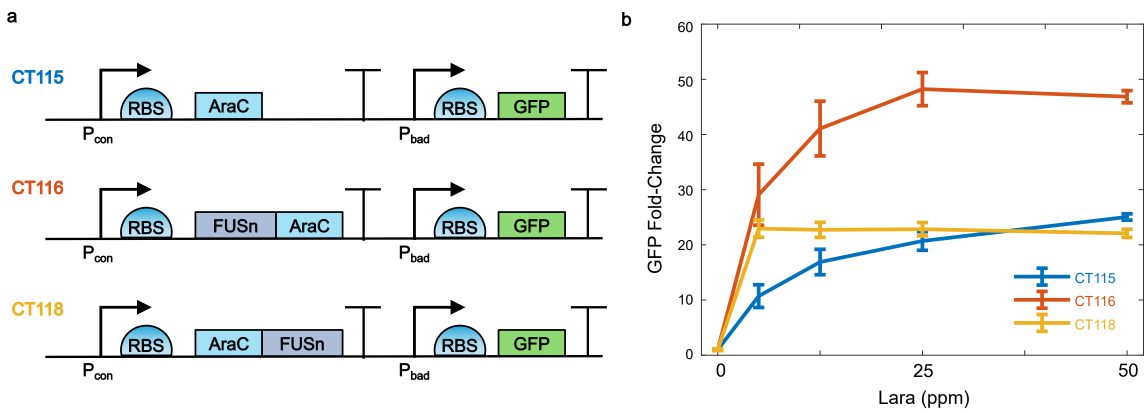


**Supplementary figure 4. Circuit Designs and Dose-Response Analysis of AraC-FUSn Fusions in open-loop circuits.** (a) Schematic representation of the CT115, CT116, and CT118 circuit designs. In these open-loop constructs, AraC is either expressed alone or fused to the N- or C-terminus of FUSn, all under a constitutive promoter. The reporter gene GFP is controlled by the Pbad promoter. (b) Dose-response curves illustrating GFP fold-changes as a function of inducer concentration for circuits CT115, CT116, and CT118 at 37°C. Data are displayed as mean ±s.d. (n=3).

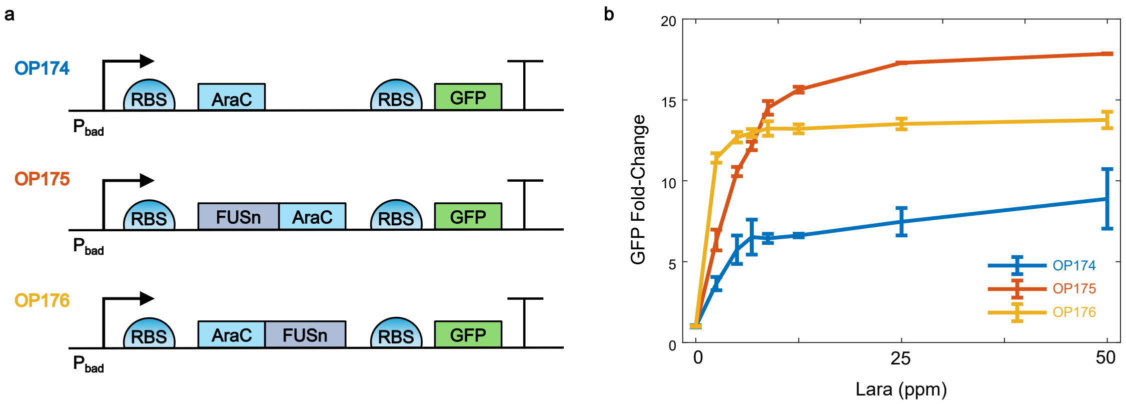


**Supplementary figure 5. Circuit Designs and Dose-Response Analysis of AraC-FUSn Fusions in closed-loop SA circuits.** (a) Schematic representation of OP174, OP175 and OP176 circuit designs. Here, AraC is either expressed alone or fused to the N- or C-terminus of FUSn under control of Pbad promoter along with the reporter gene GFP. (b) Dose-response curves illustrating GFP fold-changes as a function of inducer concentration for circuits OP174, OP175 and OP176 at 37°C. Data are displayed as mean ±s.d. (n=3).

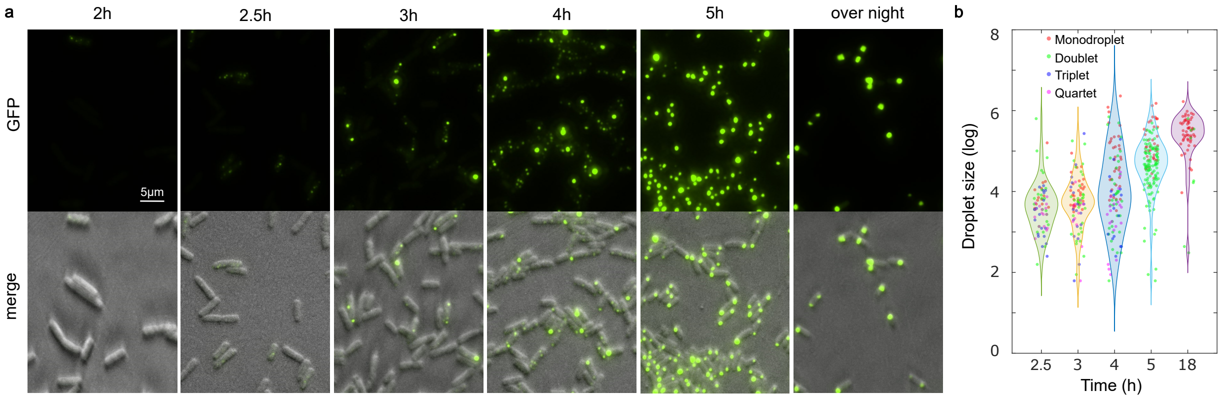


**Supplementary figure 6. Dynamics of Droplet Formation in Cells Expressing GFP-RLP20-AraC (OP203).** (a) Fluorescence microscopy images depicting the dynamics of droplet formation in cells induced to express GFP-RLP20-AraC (OP203) for various time points: 2, 2.5, 3, 4, and 5 hours, as well as overnight, with 50 ppm Lara at 34°C. (b) Quantification of droplet size and number in individual cells at different time points. Each dot represents a single droplet.

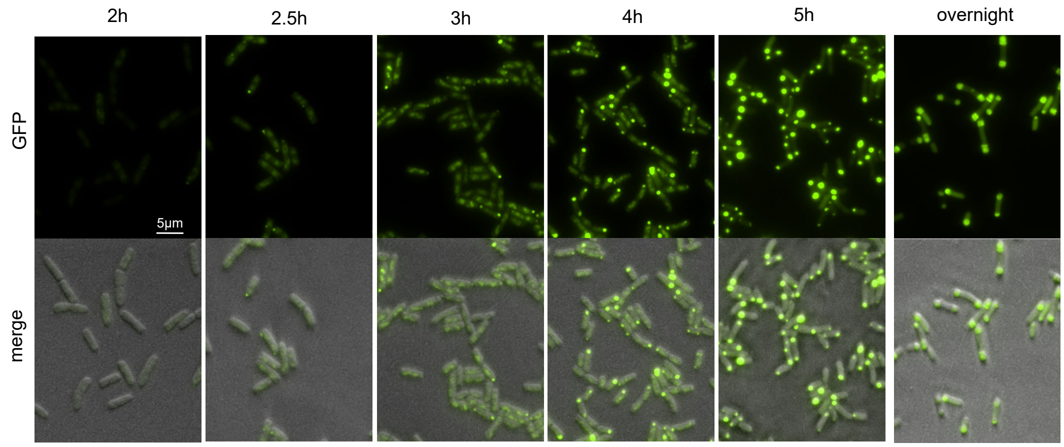


**Supplementary figure 7. Dynamics of Induced Expression of GFP-GSlinker-AraC (OP177).** Fluorescence microscopy images illustrating the dynamics of induced expression of the fusion protein GFP-GSlinker-AraC (OP177) over time: 2, 2.5, 3, 4, and 5 hours, as well as overnight, following induction with 50 ppm Lara at 34°C.

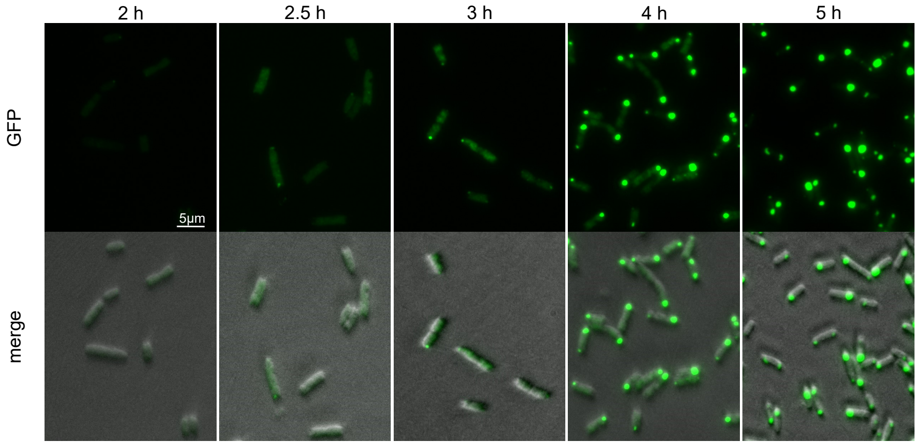


**Supplementary figure 8. Dynamics of Induced Expression of GFP-AraC (OP152).** Fluorescence microscopy images illustrating the dynamics of induced expression of the fusion protein GFP-AraC (OP152) over time: 2, 2.5, 3, 4, and 5 hours following induction with 50 ppm Lara at 34°C.

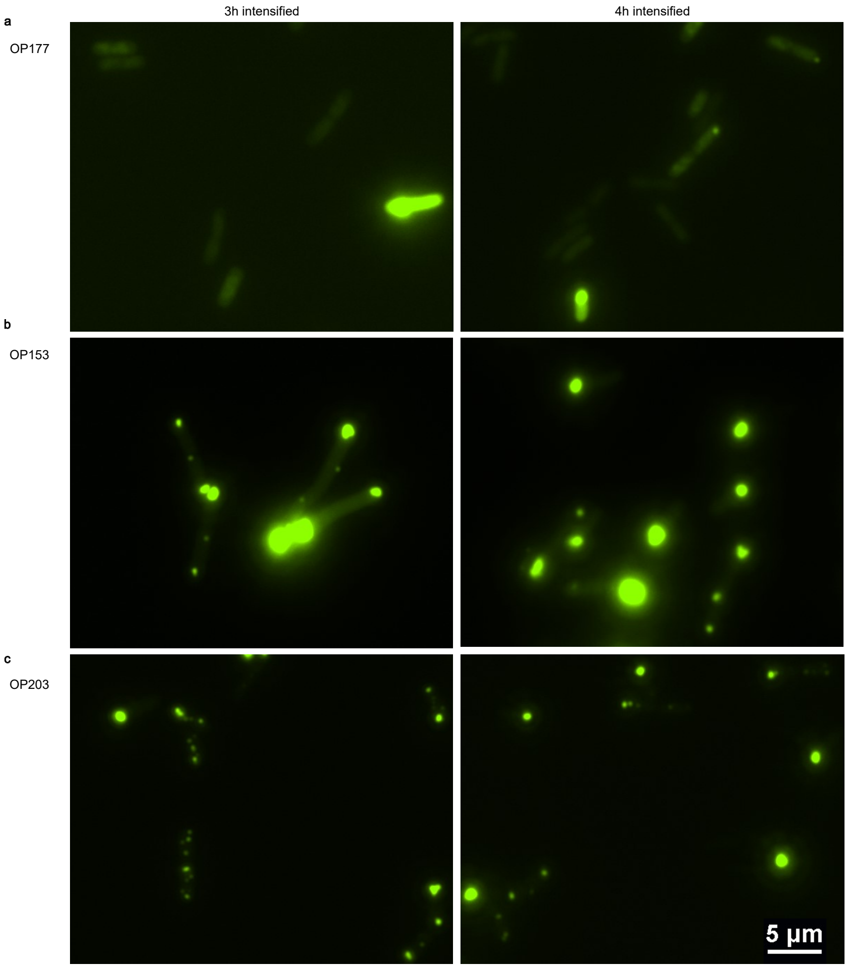


**Supplementary figure 9.** Intensified fluorescence microscopy images revealed that cells containing circuits OP153 and OP203 maintained small droplets at 3~4 hours post-dilution, whereas cells with circuit OP177 exhibited a notable absence of GFP in most cells. These images are the same as those shown in Fig. 4a-c but with intensified fluorescence.

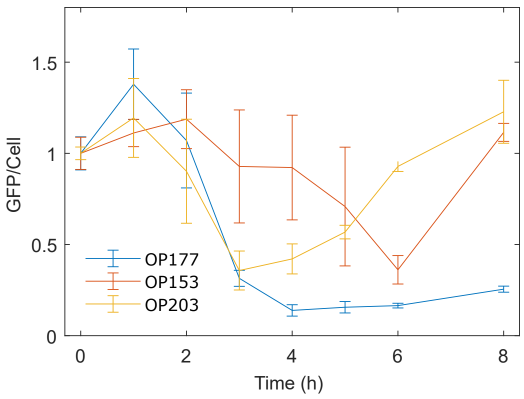


**Supplementary figure 10.** Time-course analysis of average GFP levels per cell following the dilution of cells expressing the activated OP177, OP153, and OP203 circuits into the fresh medium with 2.5 ppm Lara ±at 34°C. Data are presented as means ± s.d. (n ≥ 3. n represents number of images for each data point.) Total fluorescence intensity of all the cells in each image was measure by ImageJ with background fluorescence being deducted.

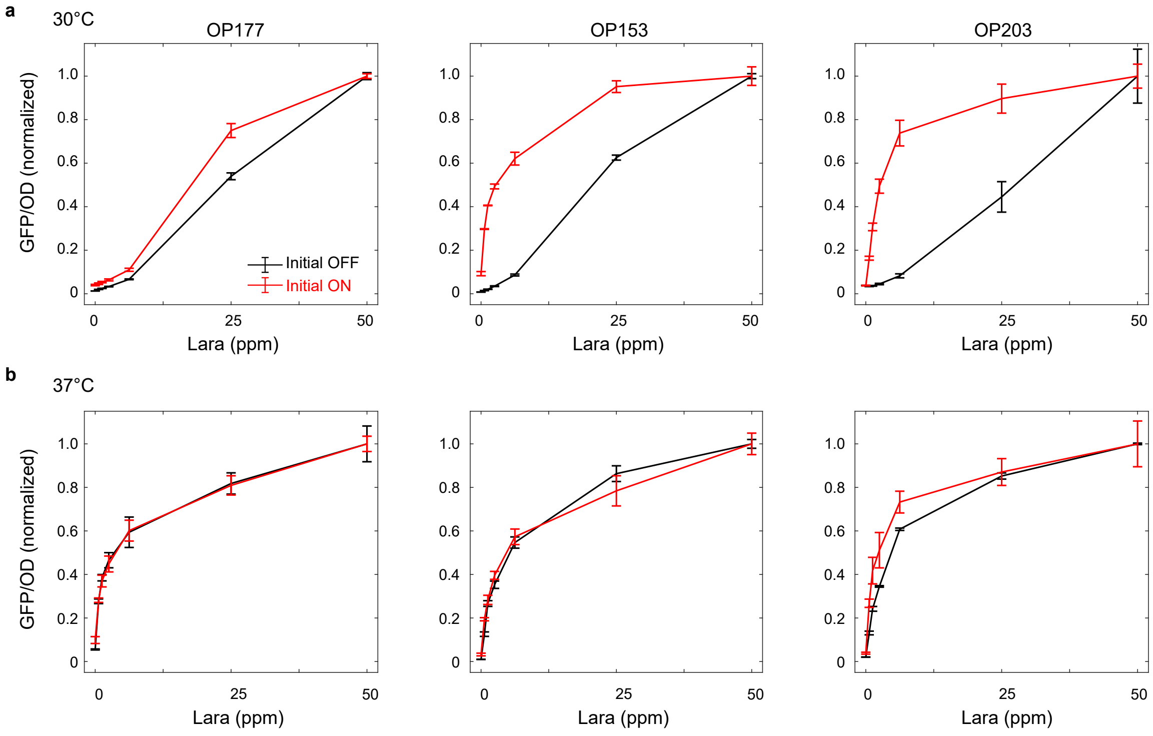


**Supplementary figure 11.** Hysteresis analysis of circuits OP177, OP153, or OP203 at Different Temperatures. Dose-response curves illustrate the steady-state GFP/OD levels in cells containing circuits OP177 (left), OP153 (middle), or OP203 (right), starting from either OFF or ON states at 30°C (a) or 37°C (b). Data are presented as mean ± s.d. (n = 3).

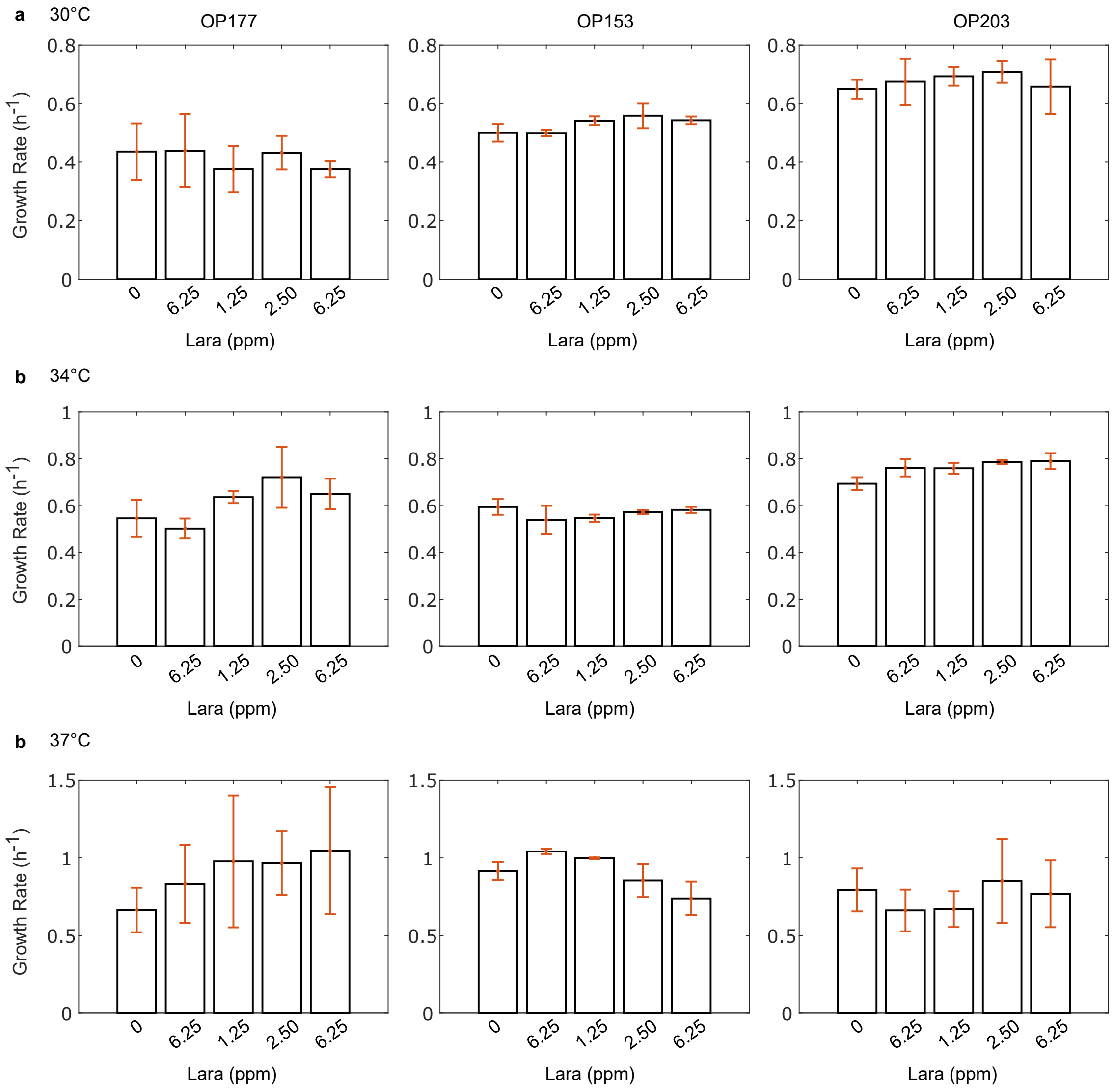


**Supplementary figure 12.** Growth rates of cells expressing activated circuits OP177, OP153, or OP203 at temperatures of 30°C (a), 34°C (b), and 37°C (c) with different concentrations of Lara after dilution. Data are presented as means ± s.d. (n = 3).

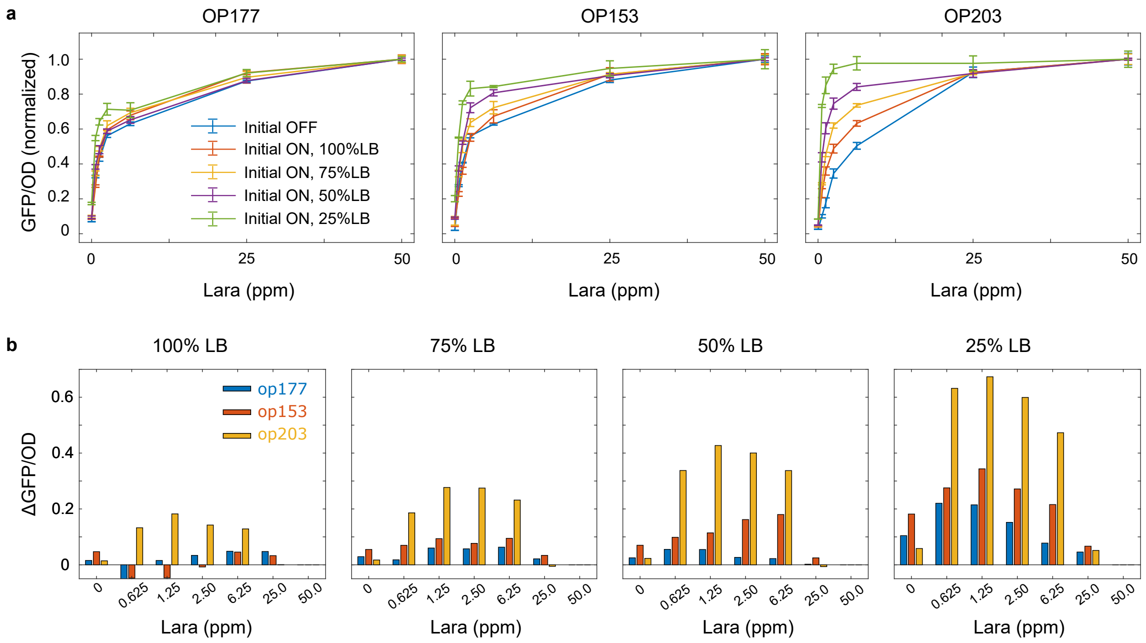


**Supplementary figure 13. Dose-Response Analysis of Steady-State GFP/OD Levels Under Varying Nutrient Conditions.** (a) Dose-response curves illustrating steady-state GFP/OD levels in cells containing circuits OP177, OP153, or OP203, grown under different nutrient levels, starting from either OFF or ON states at 37°C. Data are presented as means ± s.d. (n = 3). (b) Comparison of steady-state GFP/OD levels between cells starting from ON versus OFF states (ΔGFP/OD) in each system under varying nutrient levels. Different percentages of LB media were prepared by mixing certain amounts of LB broth with M9 salt. For example, 75 % LB was made by mixing 75 parts of LB broth with 25 parts of M9 salt.
